# Constructing a Draft Map of the Cannabis Proteome

**DOI:** 10.1101/577635

**Authors:** Conor Jenkins, Ben Orsburn

**Affiliations:** Think20 Labs, Columbia, MD; Hood College Department of Biology, Frederick, MD; Proteomics und Genomics, Baltimore, MD

## Abstract

Recently we have seen a relaxing on the historic restrictions on the use and subsequent research on the cannabis plants, generally classified as *Cannabis sativa* and *Cannabis indica*. What research has been performed to date has centered on chemical analysis of plant flower products, namely cannabinoids and various terpenes that directly contribute to phenotypic characteristics of the female flowers. In addition, we have seen many groups recently completing genetic profiles of various plants of commercial value. To date, no comprehensive attempt has been made to profile the proteomes of these plants. In this study we present our initial findings consisting of the identification of 17,269 unique proteins identified from Cannabis plant materials, as well as 6,110 post-translational modifications identified on these proteins. The results presented demonstrate the first steps toward constructing a complete draft map of the Cannabis proteome.

## Introduction

Proteomics is a science dedicated to the creation of comprehensive quantitative snapshots of all the proteins produced by an individual organism, tissue or cell.^1^ The term was coined in the 1990s during the efforts to sequence the first complete human genomes.^2^ While the technology was in place to complete the human genome draft in 2003, the first two drafts of the human proteome were not completed by teams led by Johns Hopkins and CIPSM researchers until 2014. These two separate and ambitious projects were the composite of thousands of hours of instrument run time on the most sophisticated hardware available at that time.^3,4^ Recent advances in mass spectrometry technology now permit the completion of proteome profiles in more practical time. Single celled organisms have been “fully” sequenced in less than an hour, and by use of multi-dimensional chromatography, relatively high coverage human proteomes have been completed in only a few days.^5–7^ While much can be learned by sequencing DNA and RNA in a cell, quantifying and sequencing the proteome has distinct advantages as proteins perform physical and enzymatic activities in the cell and are therefore more directly linked to cell physical characteristics.^8^ Furthermore, many proteins are altered by chemical post-translational modifications such as phosphorylation and acetylation which may completely change the protein function by serving as on/off switches for motion or metabolism.^9,10^ RNA sequencing may correctly predict the presence and relative abundance of proteins, but proteins inactivated by chemical modifications may make predictions of function from RNA abundance data wholly inaccurate. Protein modifications are directly involved in nearly every known disease and these modifications are impossible to identify with any current DNA/RNA sequencing technology.^11^

In North America we have recently witnessed the relaxing of historic restrictions on the use and subsequently, the research, of the plants of the *Cannabis* genus. To date, relatively little work has been performed on these plants in any regard and no comprehensive study of the proteome has ever been attempted. Table 1 describes the three studies performed to date on these plants. In a study published in 2004, Reharjo *et al.*, described a differential proteomics approach for the studying of *Cannabis sativa* plant tissues. Differential analysis was performed by two-dimensional gel electrophoresis (2D-gel), followed by mass spectrometry. The counting of gel spots indicated at least 800 proteins were present in these tissue, but due to technological restraints of the times, less than 100 were identified.^12^ We report herein the methodology and preliminary results in our attempts to create the first draft map of the proteomes of *Cannabis*. Protein was extracted from plant tissue from stems and leaves of plants as well as from medical flower products from *C.sativa* and *C.indica* strains with well characterized cannabinoid profiles. Extracted proteins were digested, separated by ultrahigh pressure liquid chromatography coupled to ultrahigh resolution tandem mass spectrometry. Data assembly and annotation is ongoing, but we have obtained unique protein identifications corresponding to 17,269 proteins or theoretical gene products, as well as the localization of 6,110 post-translational modifications on these proteins.

**Table 1.** A summary of Cannabis Proteomic Studies

| Study | Date | Number of Proteins Identified |
| --- | --- | --- |
| Raharjo et al., | 2004 | 54 |
| Aiello et al., | 2016 | 181 |
| Vincent et al., | 2019 | 160 |
| This study |  | 17,269 |

**Figure 1).**
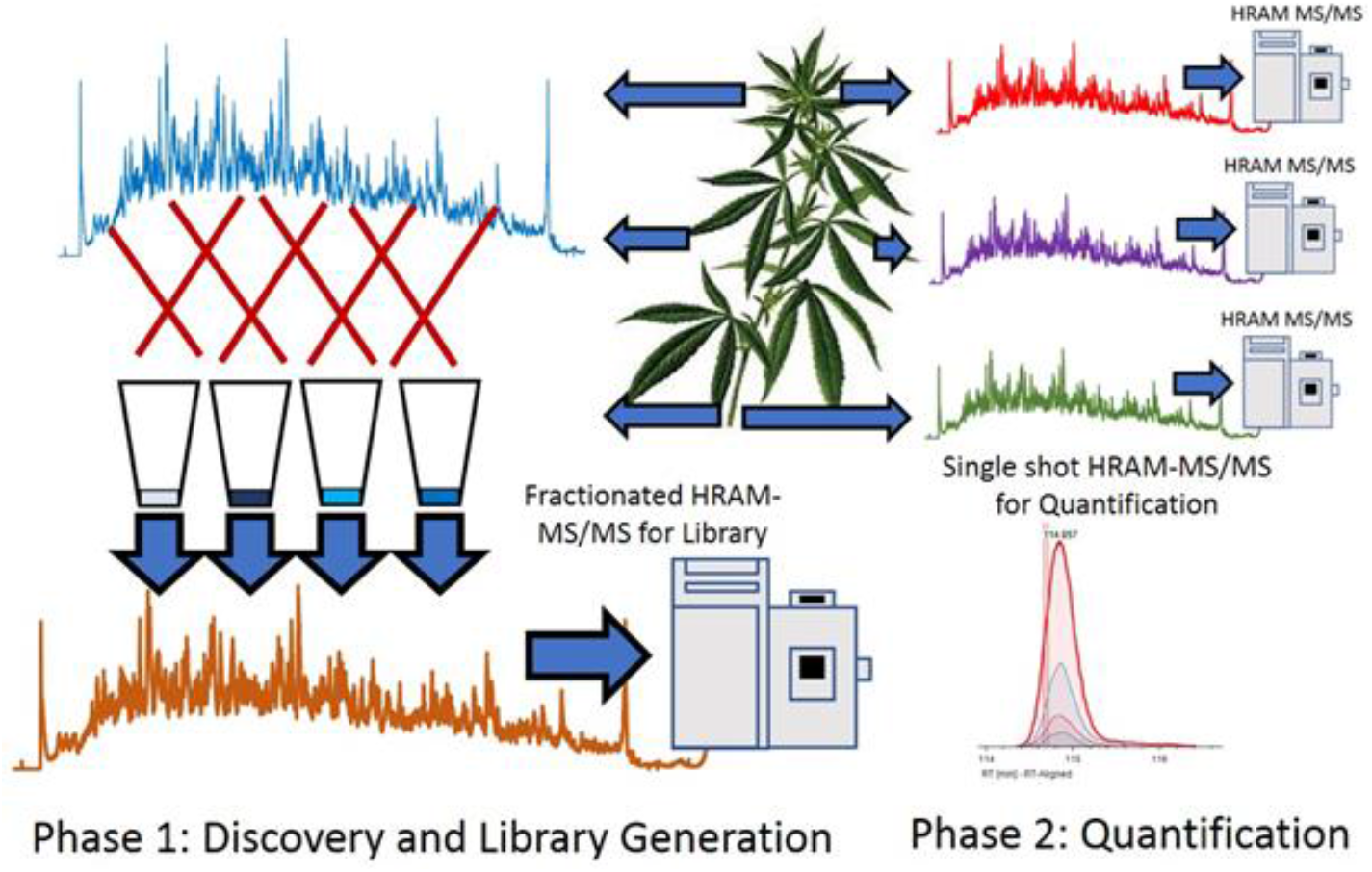
An overview of the proteomics strategy. Combined material was heavily fractionated in order to develop library data. Individual materials were separately analyzed by single-shot analysis for quantification.

## Materials and Methods

### Samples

A table of samples analyzed to date are described in Supplemental Table 1. All samples were obtained by Think20Labs under the guidelines of the MMCC regulations in accordance with a temporary license granted under COMAR 10.62.33.^13^ A recent study described the optimization of digestion conditions for the proteomic analysis of Cannabis flowers and performed similar experiments as the ones described here. Vendor instrument files are available at the MASSIVE data repository as MSV00083191.

### Sample Preparation

Multiple variations on protein extraction and digestion were tested, based on the highest percent recovery of peptides per milligram of starting plant material by use of an absorbance based assay for tryptic peptides (Pierce) (data not shown). The final sample method was based on the filter-aided sample preparation method (FASP).^14^ Briefly, 1mg of fresh plant material was flash frozen at −80°C for 20 minutes. The cell walls were disrupted by immediately removing the frozen material and blunt physical concussion. The material was then dissolved in a solution of 150 μL of 5% SDS and 0.2%DTT and heated at 95°C for 10 minutes to reduce and linearize proteins and reduced to room temperature on ice. 150 μL of 8M urea/50mM TrisHCl was added to the mixture. Detergent removal, protein cysteine alkylation and sample cleanup for digestion was performed according to the FASP protocol. All reagents were obtained from Expedeon BioSciences. Proteins were digested for 16 hours at room temperature. Digested peptides were released by centrifuging the FASP chamber at 13,000 × g for 10 minutes with peptides eluting into a new 1.5 mL centrifuge tube. An additional 75 μL of the digestion solution was added and the elution was repeated. The peptides were dried by vacuum centrifugation (SpeedVac). Peptides were resuspended in 20μL of 0.1% trifluoroacetic acid for either desalting or for high pH reversed phase fractionation. Peptides were quantified by absorbance using a peptide specific kit (Pierce).

### Peptide Fractionation

Aliquots approximating 50 micrograms of alkylated peptides were combined from each sample into a shared pool for offline fractionation and library generation. Due to sample availability constraints, two batches containing peptides from 6 separate samples were fractionated separately. High resolution fractionation followed a recent protocol^7^, with the exception that an Accela pump (Thermo) was utilized for gradient delivery and fractions were concatenated following collection. The concatenation was performed as described previously.^15^

### LC-Mass Spectrometry Analysis

For initial analysis all fractionated and single shot samples were ran identically on a nanoESI-Q Exactive HF-X system. Briefly, 2 μg of peptides were loaded into a 4cm trap column and eluted with an optimized gradient on a 100 cm monolithic 75μm column utilizing a 135 minute total elution gradient. Eluting peptide masses were acquired at 120,000 resolution followed by the fragmentation of the most abundant eluting peptides with HCD fragmentation at 27eV. Fragmented peptides were acquired at 15,000. Although this system is capable of higher scan speed, a higher resolution MS/MS was utilized in order to obtain more confident identification and localization of PTMs. The top 15 most abundant ions were selected for fragmentation. Each single shot sample injected twice, once with a 30ms ion injection time, and again with a 150ms ion injection time. Dynamic exclusion was utilized allowing each ion to be fragmented once, any ion within 5ppm of the matched ion was excluded from fragmentation for 60 seconds, or approximately 2.2x the peak width.

### Peptide and Protein Identification

An overview of the data processing pipeline is shown in Figure 2. To date, no full annotated protein FASTA exists for any Cannabis species. Classical proteomics workflows require a reference theoretical protein database from which to construct matches from MS1 and MS/MS spectral data. In lieu of this we utilized 2 sources of information for identifying MS/MS spectra. As less than 600 annotated sequences for Cannabis exist in the UniProt library, a custom UniProt/SwissProt database consisting of every manually annotated sequence from green plants was used. In addition, three high quality genome sequences publicly available were subjected to 6-frame translation in house using the MaxQuant plug-in^16^, to create theoretical protein sequences that accurately match the material being analyzed. These files are described in Figure 2 and details are listed in Supplemental Table 5. For initial analysis, all data processing was performed in Proteome Discoverer 2.2 (PD) (Thermo Fisher) using the SequestHT, Percolator and Minora algorithms. The proteogenomic FASTA was crudely reduced during database import in PD according to manufacturer default settings. SequestHT and Percolator produce identified and confidence scored peptide spectral matches (PSMs). Multiple consensus workflows were used within PD to assemble the PSMs into peptide groups, protein database matches, and finally non-redundant proteins groups using the principle of strict parsimony as defined by the vendor software defaults. Figure 2 describes the sequence databases employed in this study. All settings used in PD for peptide spectral match and protein identification are described in Supplemental Table 2.

**Figure 2).**
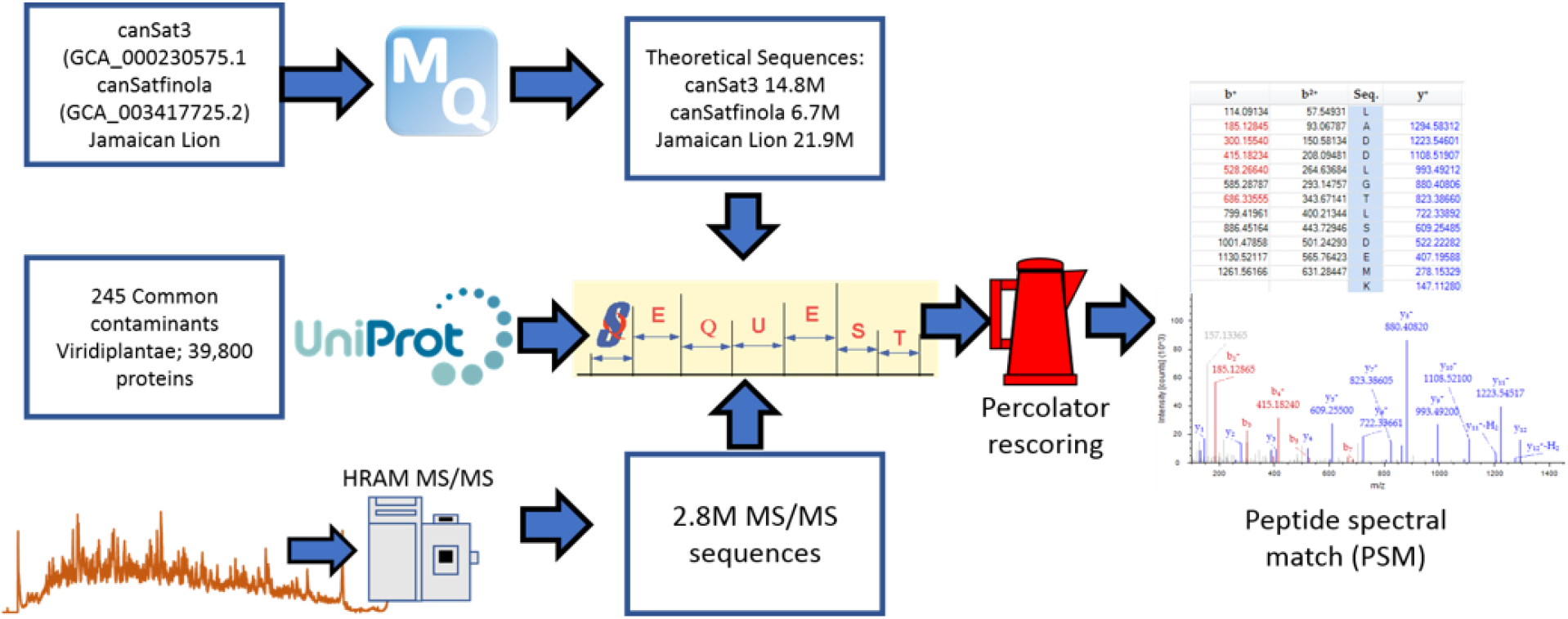
An overview of the data processing pipeline inputs and algorithms used for initial analysis and generation of peptide spectral matches.

**Figure 3.**
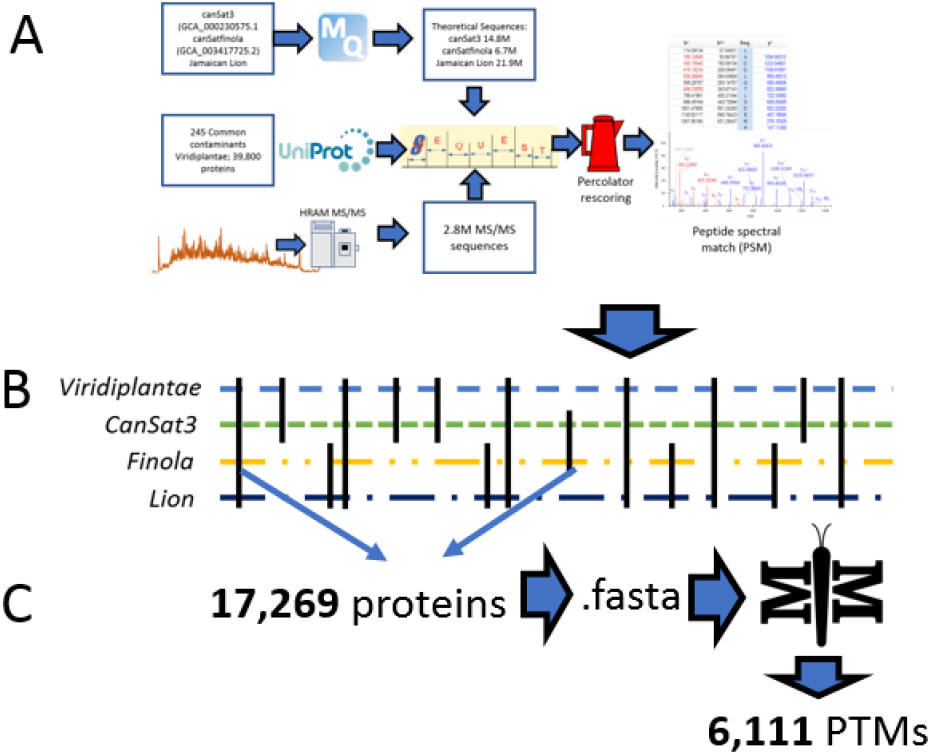
**A)** PSMs matched to all databases described. **B)** Identified PSMs match 58,309 protein sequences that can be reduced to 17,269 unique protein groups. **C)** This list of conserved proteins can be used to create a smaller database that can be used by MetaMorpheus to search all MS/MS spectra for PTMs

### Chromosome alignment

A recent re-analysis of the CanSat3 genome^17^ aligned the sequences into ten separate chromosome files.^18^ The Protein Marker node in PD was used in 4 rounds of reprocessing of the consensus processing to develop a metric of the number of identified protein entries in this study that are products of each chromosome. Four rounds were necessary due to a limitation in the software that allows a maximum of 3 separate FASTA sequences to be used for output marking. Reiterations of this analysis were repeated to ensure that the chromosomes grouped in each re-analysis was an independent variable and did not affect localization output (data not shown).

Both protein data, representing potential redundancies and unique protein groups were obtained. The results are plotted in Figure 5.

### Identification of PTMs

The recently described MetaMorpheus software package (MM) was used for the indiscriminate identification of post-translational modifications. The full theoretical protein sequences of the 17,269 unique proteins obtained from the PD analysis were extracted in FASTA format and used for MM analysis using the default workflows for Recalibration, GPTMD, Search and Post Processing.^19,20^

### Graph generation

The UpSetR package was used for the comparison of proteome to genome sequencing files using both the webhosted ShinyApp (https://gehlenborglab.shinyapps.io/upsetr/) as well as the full package within RStudio 1.0.143. Supplemental figures were generated in the PNNL Venn Diagram 1.5.5 tool as well as with GGPlot2^21^ within RStudio.

## Results and Conclusion

### Peptide and protein identifications

The lack of extensive pre-existing information into Cannabis and the genes/proteins active is a considerable challenge for traditional proteomics workflows which rely heavily on annotated and reviewed protein sequence databases for spectral matching. Using the custom proteogenomic workflow described here, we were able to identify a small percentage of the first 2.5 million MS/MS spectra obtained and match those to a compiled and in-house generated theoretical protein sequence database of greater than 41 million entries. This pipeline resulted in 135,845 peptide spectral matches, or approximately 5.4% identification rate.

Further improvements on the Cannabis proteome are underway and will be described elsewhere. A primary focus will be the refinement and annotation of the currently existing Cannabis genomes available. Recent work has described the improvement and correction of genome annotations using high resolution mass spectrometry.^22^ While this is beyond the scope of this study, we can develop metrics related to the quality of match of genomic data using high coverage proteomics. An UpSetR graph^23^ is shown in Figure 4 that shows the unique protein identifications and matches to the various genomic databases both unique and conserved. To further illustrate the importance of an annotated Cannabis protein database, of the protein groups identified by the initial analysis, only 1,838 -- representing less than 10% of all matches had sequence homology suitable to be matched against the UniProt/SwissProt database. This is despite the fact a database containing the protein sequences of all green plants was utilized in this work. A summary of these results is available as Supplemental Table 2.

**Figure 4).**
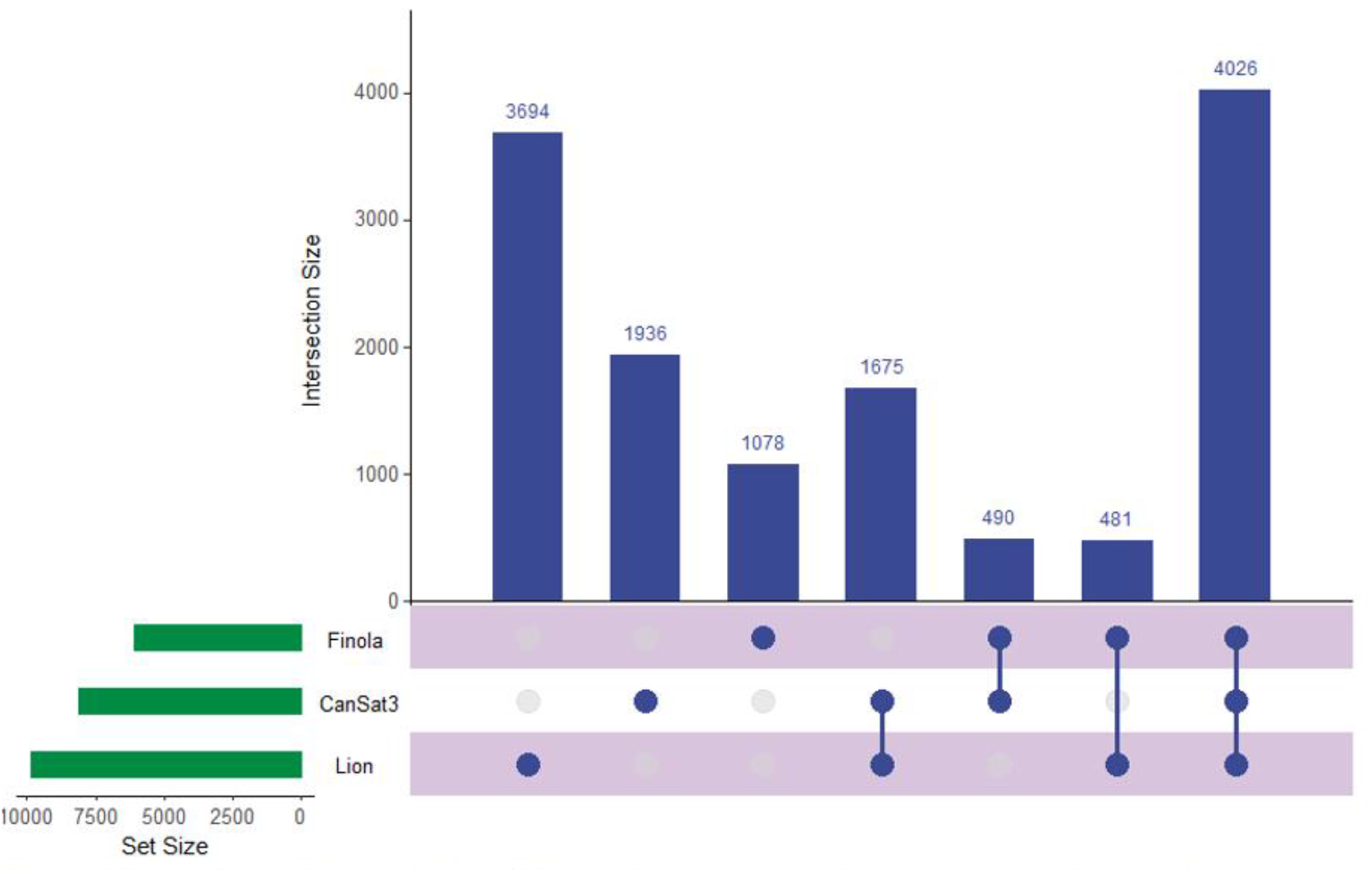
Depiction of the matches of the proteomic data to the separate next generation sequencing files utilized in this study.

**Figure 5).**
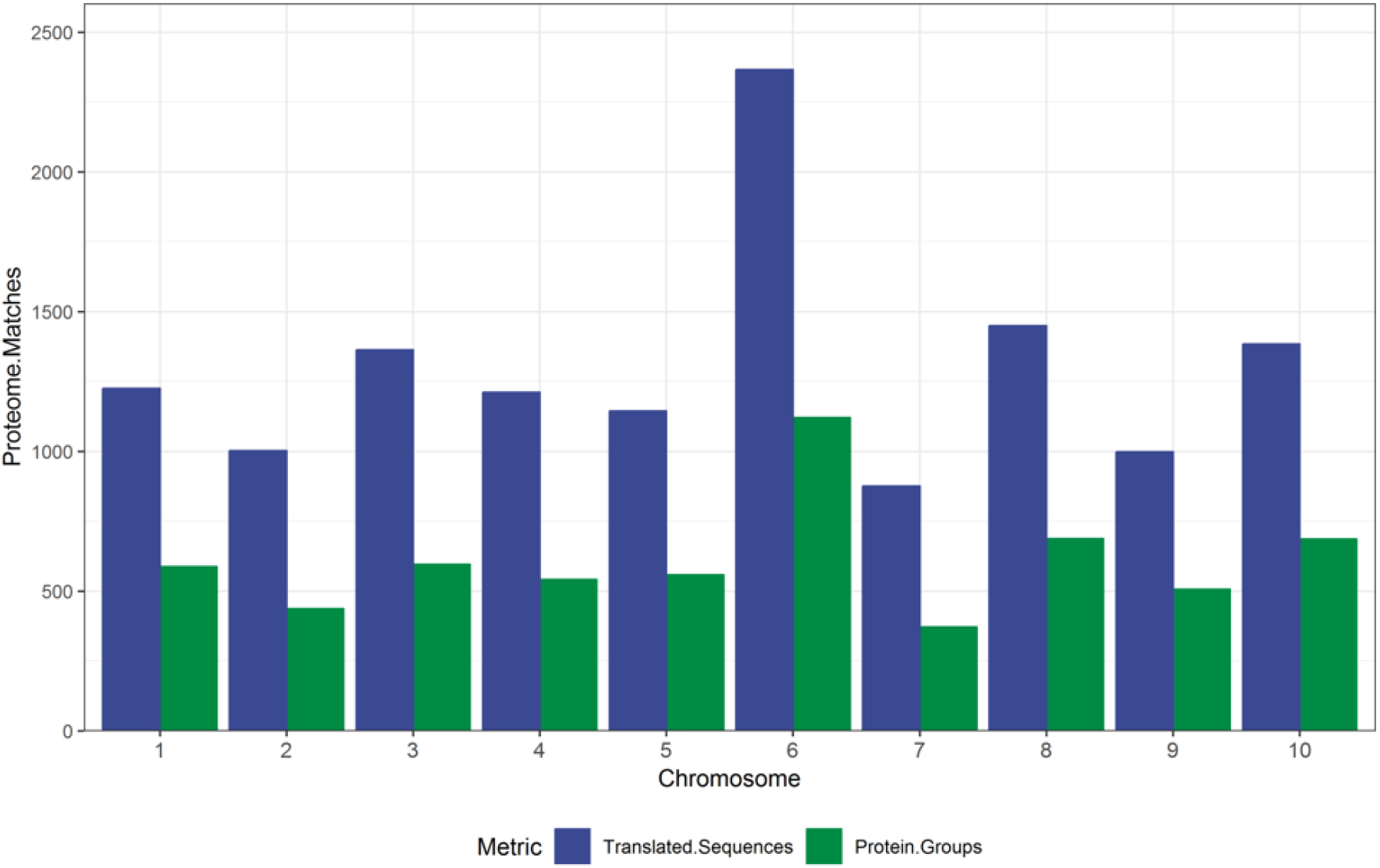
A comparison of the matches to assembled gene product and unique protein groups to the assembled chromosomes from the recent re-analysis of the CanSat3 genome.

Despite these challenges we have built considerably on the existing knowledge of the Cannabis proteome and have reason to believe that with proper annotation we have generated a relatively complete picture of the protein composition of the plants studied to date. Genomic analysis has identified approximately 25,000 genes, a number comparable to that of humans in a Cannabis chromosome assembly^24^. While exact numbers are still currently debated, no study to our knowledge as identified 20,000 or more unique protein groups.^25^

### Post-translational Modification Identification and Analysis

The importance of protein post-translational modifications in Cannabis are, to our knowledge, completely unknown. Current strategies for identifying PTMs from shotgun proteomic data require the addition of dynamic modification. Each single dynamic modification results in a doubling of the number of theoretical peptides and due to the presence of multiple modifications sites per protein, indiscriminate searching of PTMs results in exponential increase in both the search space and required computational power to complete data processing.^26^ To address these issues we generated a new FASTA database that contained only the 17,269 proteins identified in our SequestHT and Percolator searches of all high resolution files. Using this newly reduced database of proteins that appear to be present and the complete theoretical sequences from these entries extracted from our original FASTAs, we can search these identified proteins for PTMs. For this analysis we chose to employ the recently described MetaMorpheus (MM). MM performs a tiered search strategy that is reliant on the recalibration of MS spectra and the GPTMD algorithm.^27^ This next generation search engine is capable of searching for, identifying, and quantifying hundreds of unknown post-translational modifications with annotated databases on standard desktop computers.^19,20^ To test the capabilities of this workflow, we used MM to search the single shot analysis of 6 commercial flowers performed in duplicate as described. Supplemental Table 4 contains a summary of these results. MM identified 26,477 unique peptides and 6,111 PTMs in these files alone. Further analysis and validation will be necessary to determine the function and importance of these modifications in these plants.

## Conclusions

We have performed the first comprehensive proteomic analysis of Cannabis plants toward our goal of developing a draft map of the Cannabis proteome. From the samples analyzed to date and described herein, we have identified a possible 17,269 unique protein group identifications by matching these high resolution and accuracy MS/MS spectra to currently available protein and genetic sequence data available. Toward these aims, we developed a robust sample prep protocol capable of extracting peptides from flowers, leaves and stems of Cannabis plants with high efficiency as well as a bioinformatic pipeline capable of identifying both proteins and post translational modifications in these materials.

Further work is currently underway to improve upon these early data with emphasis on the bioinformatic workflows necessary to annotate and interpret these data.

## Acknowledgements

We’d like to thank Edward Sawicki for his insight and helpful discussions in the construction of this study and manuscript.

