## Supplemental 2 for "Constructing a Draft Map of the Cannabis Proteome"

------------------------------------------------------------------

The pipeline tree:

------------------------------------------------------------------

|-(0) Spectrum Files

|-(1) Spectrum Selector

|-(3) Sequest HT

|-(4) Percolator

|-(2) Minora Feature Detector

------------------------------------------------------------------

Processing node 0: Spectrum Files

------------------------------------------------------------------

------------------------------------------------------------------

Processing node 1: Spectrum Selector

------------------------------------------------------------------

1. General Settings:

- Precursor Selection: Use MS1 Precursor

- Use New Precursor Reevaluation: True

- Use Isotope Pattern in Precursor Reevaluation: True

2. Spectrum Properties Filter:

- Lower RT Limit: 0

- Upper RT Limit: 0

- First Scan: 0

- Last Scan: 0

- Lowest Charge State: 0

- Highest Charge State: 0

- Min. Precursor Mass: 350 Da

- Max. Precursor Mass: 5000 Da

- Total Intensity Threshold: 0

- Minimum Peak Count: 1

3. Scan Event Filters:

- MS Order: Is Not MS1

- Min. Collision Energy: 0

- Max. Collision Energy: 1000

- Scan Type: Is Full

4. Peak Filters:

- S/N Threshold (FT-only): 1.5

5. Replacements for Unrecognized Properties:

- Unrecognized Charge Replacements: Automatic

- Unrecognized Mass Analyzer Replacements: ITMS

- Unrecognized MS Order Replacements: MS2

- Unrecognized Activation Type Replacements: CID

- Unrecognized Polarity Replacements: +

- Unrecognized MS Resolution@200 Replacements: 60000

- Unrecognized MSn Resolution@200 Replacements: 30000

6. Precursor Pattern Extraction:

- Precursor Clipping Range Before: 2.5 Da

- Precursor Clipping Range After: 5.5 Da

------------------------------------------------------------------

Processing node 3: Sequest HT

------------------------------------------------------------------

1. Input Data:

- Protein Database:

canSat3_genome.out.fasta

contaminants.fasta

finola1.out.fasta

Jamaican_Lion_polished_primary_181018.out.fasta

viridiplantae_reviewed_fasta.fasta

- Enzyme Name: Trypsin (Full)

- Max. Missed Cleavage Sites: 2

- Min. Peptide Length: 6

- Max. Peptide Length: 150

- Max. Number of Peptides Reported: 10

2. Tolerances:

- Precursor Mass Tolerance: 10 ppm

- Fragment Mass Tolerance: 0.02 Da

- Use Average Precursor Mass: False

- Use Average Fragment Mass: False

3. Spectrum Matching:

- Use Neutral Loss a Ions: True

- Use Neutral Loss b Ions: True

- Use Neutral Loss y Ions: True

- Use Flanking Ions: True

- Weight of a Ions: 0

- Weight of b Ions: 1

- Weight of c Ions: 0

- Weight of x Ions: 0

- Weight of y Ions: 1

- Weight of z Ions: 0

4. Dynamic Modifications:

- Max. Equal Modifications Per Peptide: 3

- Max. Dynamic Modifications Per Peptide: 4

7. Static Modifications:

- 1. Static Modification: Carbamidomethyl / +57.021 Da (C)

------------------------------------------------------------------

Processing node 4: Percolator

------------------------------------------------------------------

1. Input Data:

- Maximum Delta Cn: 0.05

- Maximum Rank: 0

2. Decoy Database Search:

- Target FDR (Strict): 0.01

- Target FDR (Relaxed): 0.05

- Validation based on: q-Value

------------------------------------------------------------------

Processing node 2: Minora Feature Detector

------------------------------------------------------------------

1. Peak & Feature Detection:

- Min. Trace Length: 5

- Min. # Isotopes: 2 Peaks

- Max. ΔRT of Isotope Pattern Multiplets [min]: 0.2

2. Feature to ID Linking:

- PSM Confidence At Least: High
